# Understanding the role of apolipoproteinA-I in atherosclerosis. Post-translational modifications synergize dysfunction?

**DOI:** 10.1101/2020.06.09.142844

**Authors:** Ivo Díaz Ludovico, Romina A. Gisonno, Marina C. Gonzalez, Horacio A. Garda, Nahuel A. Ramella, M. Alejandra Tricerri

## Abstract

**Background:** The identification of dysfunctional human apolipoprotein A-I (apoA-I) in atherosclerotic plaques suggests that protein structure and function may be hampered under a chronic pro inflammatory scenario. Moreover, the fact that natural mutants of this protein elicit severe cardiovascular diseases (CVD) strongly indicates that the native folding could shift due to the mutation, yielding a structure more prone to misfold or misfunction. To understand the events that determine the failure of apoA-I structural flexibility to fulfill its protective role, we took advantage of the study of a natural variant with a deletion of the residue lysine 107 (K107del) associated with atherosclerosis.

**Methods:** Biophysical approaches, such as electrophoresis, fluorescence and spectroscopy were used to characterize proteins structure and function, either in the native conformation or under oxidation or intramolecular crosslinking.

**Results:** K107del structure was more flexible than the protein with the native sequence (Wt) but interactions with artificial membranes were preserved. Instead, structural restrictions by intramolecular crosslinking impaired the Wt and K107del lipid solubilization function. In addition, controlled oxidation decreased the yield of the native dimer conformation for both variants.

**Conclusions:** We conclude that even though mutations may alter protein structure and spatial arrangement, the highly flexible conformation compensates the mild shift from the native folding. Instead, post translational apoA-I modifications (probably chronic and progressive) are required to raise a protein conformation with significant loss of function and increased aggregation tendency.

**General Significance:** The results learnt from this variant strength a close association between amyloidosis and atherosclerosis.

**Graphical abstract:** 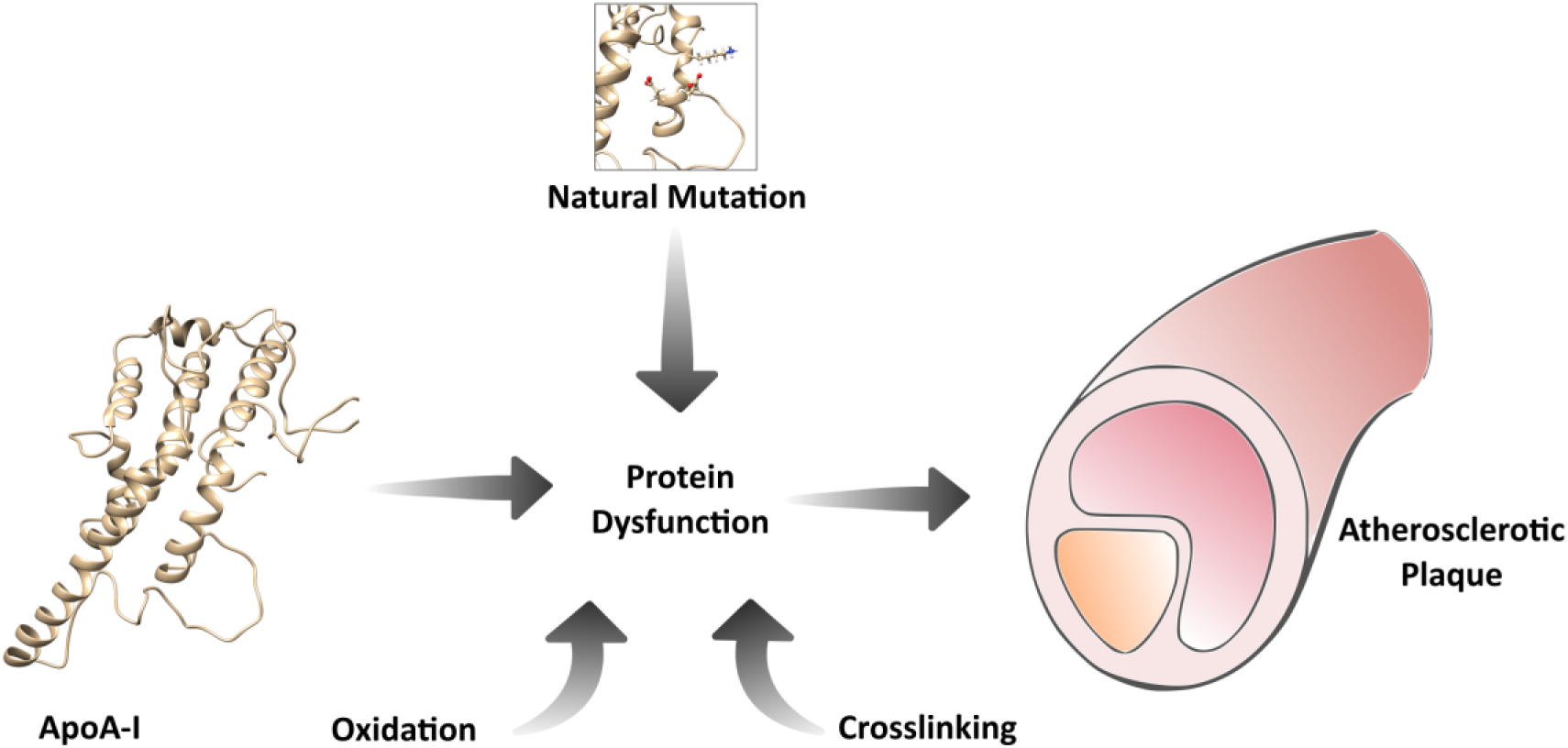

**Highlights:**

-Oxidation is clue to induce protein misfolding
-Natural mutation does not seem critical as a sole reason to determine pathogenicity
-Atherosclerosis and amyloidosis are closely related
-Intramolecular crosslinking restrains protein flexibility and function

## 1. Introduction

An extensive research field supports a key role of human high density lipoproteins (HDL) and their major protein apolipoprotein A-I (apoA-I) in the protection against cardiovascular disease [1][2]. Many functions may be involved in such clue events. Traditional evidence supported their beneficial participation in the reverse cholesterol transport (RCT), which delivers cholesterol back to the liver for its catabolism [3]. In addition, it has been recognized a crucial participation of HDL and apoA-I in pathways protecting endothelial functions: stimulation of nitric oxide-mediated vasodilatation [4], reduction of vascular cell adhesion molecule-1 (VCAM-I) expression [5][6] and the inhibition of apoptosis promoting proliferation [7]. The multiple functions of apoA-I have been extensively studied and related to a highly flexible structure. A dynamic inter conversion between lipid-free and lipid bound states mediates its efficiency to interact with membranes and to recruit phospholipids and cholesterol. The most of the protein circulates bound as complex HDL particles, and a minor fraction, about 5 % is recycled as the lipoprotein is catabolized, yielding lipid-free or lipid-poor conformations which are more effective to interact with key proteins such as ATP-binding cassette proteins A1 and G1 (ABCA1 and ABG1 receptors) and lecithin:cholesterol acyltransferase (LCAT) ([8] and references described there). Moreover, many other functions of apoA-I have been identified *in vitro*, suggesting that the protein mimics some of the pathways observed for HDL, as decreasing the respiratory burst induced by oxidized low-density lipoproteins (oxLDL) [9] and binding to lipopolysaccharide (LPS) [10][11].

In spite of the previous information, the “HDL-cholesterol hypothesis” has been revised as it seemed to be oversimplified by setting an atherosclerosis-risk indexfrom the low versus high-density lipoproteins ratio (LDL/HDL) [12][8]. Beyond the usefulness of this ratio as a clinical parameter, atherosclerosis is a complex chronic pathological scenario, ending up with a pro-oxidant environment with the release of reactive oxygen species (ROS) related to the failure of the cholesterol homeostasis. Interestingly, atherosclerotic plaques are characterized by dysfunctional apoA-I aggregated in the artery walls [13]. The distribution of the protein in human aorta is quite distinct from the circulating conformations. It is identifiedin high amounts deposited in chronic lesions, predominantly lipid-poor, not associated with HDL, extensively oxidized and cross-linked, and functionally impaired [14]. Whether the protein misfolding is a cause or a consequence of this micro environment is not known. In this regard, it was shown the presence of diffuse deposits of apoA-I as amyloid patches in complicated atherosclerotic plaques [15]. This fact indicates that amyloidosis and atherosclerosis may be closely associated [16].

About 20 natural variants of apoA-I have been described inducing amyloidosis with deposit and failure of clue organs such as heart, liver or kidney [17]. Interestingly, one naturally occurring deletion variant (K107del) was described inducing a unique pathologic pattern, as amyloidosis was associated with severe atherosclerosis [18]. The facts that determine this behavior are far to be known. Either the loss-of -function, increased tendency to misfold or the ability to elicit a pro inflammatory environment could mediate its role in these diseases [19]. Taking advantage of the structural comparative study among K107del and the protein with the native sequence, we set up here to understand the reasons that affect apoA-I misfunction and aggregation in the atherosclerotic plaques. Deep structural knowledge of this variant may help to clarify the participation of apoA-I (or its failure to fulfill the protective roles). As oxidation and crosslinking events are present in atherosclerotic plaques, we analyzed the variant’s natural flexibility and function, under native conditions and under controlled chemical modifications that could mimic those suffered in a chronic pathological scenario.

## 2. Materials and methods

### 2.1. Materials

Guanidine hydrochloride (GndHCl), thioflavin T (ThT), cholesterol (Chol), sodium cholate, ethylenediaminetetraacetic acid (EDTA), sodium chloride (NaCl), sodium dodecyl sulfate (SDS), Terbium -III chloride (Tb) and dipicolinic acid (DPA) were from Sigma-Aldrich (St Louis, MO); 1-palmitoyl-2-oleoylphosphatidylcholine (POPC) and dimyristoyl phosphatidylcholine (DMPC) were purchased from Avanti Polar Lipids (Alabaster, AL); His-purifying resin was from Novagen (Darmstadt, Germany). Bis-(sulfosuccinimidyl) suberate (BS^3^) and isopropyl-β-D-thiogalactoside (IPTG) were purchased to Thermo Scientific (Waltham, MA). All other reagents were of the highest analytical grade available.

### 2.2. Cloning, expression and purification of wild-type and K107del apoA-I

Proteins were expressed as previously described in [20][21]. The cDNA of apoA-I with the native sequence (Wt) and K107del were modified to introduce an acid labile Asp–Pro peptide bond between amino acid residues 2 and 3 of apoA-I, which allowed specific chemical cleavage of an N-terminal His-Tag fusion peptide. These constructs, inserted into a pET-30 plasmid (Novagen, Madison, WI) were transformed into BL21 (DE) Escherichia coli cells (Novagen, Madison, WI), then expressed by induction with IPTG and purified by elution through Ni-chelating columns (Novagen, Madison, WI) as described [21], resulting in a high yield of protein with a purity of at least 95% (determined by SDS-PAGE).

### 2.3. Crosslinking of Proteins

Proteins were crosslinked at 0.05 mg/mL in PBS pH 7.9 for 3 h without agitation at room temperature. Fresh BS^3^ was added (within 1 min after solubilization in PBS to avoid hydrolysis of free crosslinker) at 30:1 BS^3^:protein molar ratio [22]. Reactions were quenched for 15 min by the addition of Tris buffer to a final concentration of 50 mM. The concentration of crosslinked proteins was calculated by the BCA Protein Assay Kit (Thermo Scientific (Waltham, MA)). The presence of the monomeric conformation after the treatments was confirmed by polyacrylamide gradient electrophoresis, either under native or denaturing conditions. In order to evaluate whether protein oxidation may induce an aggregationprone conformation, proteins were incubated by 24 h at 0.6 mg/mL and crosslinked as described above.

### 2.4. Protein oxidation

Wt and K107del were chemically oxidized (ox-) by overnight incubation in a 10,000 molar excess of H_2_O_2_ at 37°C for 12 h. The concentration of H_2_O_2_ was determined spectrophotometrically (ε 39.4 M^-1^ cm^-1^ at 240 nm) [23]. Excess of H_2_O_2_ was removed by extensive dialysis against 10 mM sodium phosphate buffer pH 6.0 (fibrillation buffer). The molecular weights (Mw) of the oxWt and oxK107del proteins were calculated by intact protein mass spectrometry analysis. This analysis was performed at the Proteomics Core Facility CEQUIBIEM, at the University of Buenos Aires/CONICET (National Research Council). Samples were analyzed using an Ultraflex II Bruker Daltonics UV-MALDI-TOF-TOF mass spectrometer. Spectra were obtained in positive linear mode (LP), within a mass range of 20,000-60,000 m/z. The generated spectra were visualized and compared with Flex Analysis 3.3 software.

### 2.5. Fluorescence measurements

Trp intrinsic fluorescence emission spectra were acquired on an Olis upgraded SLM4800 spectrofluorometer (ISS Inc, Champaign, IL). Proteins, either freshly resuspended or after chemical modifications were diluted to 0.05-0.2 mg/mL in Tris 20 mM pH 7.4 or PBS pH 7.4. Spectra were registered fixing excitation wavelength at 295 nm as previously registered [24][21].

### 2.6. Protein/lipids interaction methodologies

#### 2.6.1. Unilamellar liposomes construction

One mg of POPC or (POPC:Chol at a 4:1 molar ratio) from a stock solution in chloroform, were used to form a film in a round-bottom tube, dried by blowing a N_2_ atmosphere and exhaustively exposed to vacuum in a lyophilizer (Virtis) to evaporate the solvent. Then, the fluorescent complex [Tb-DPA] was achieved by the addition of 15 mM Tb and 150 mM DPA in 10 mM Tris–HCl pH 8.0 added to final concentration 1mg/mL of lipids. Multilamellar liposomes (MLV) were attained by extensive vortexing, followed by sonication to rearrange lipids into small unilamellar vesicles (SUVs). Excess of the fluorescent complex was removed by elution of the SUVs through a Superose 6 HR 10/30 column equilibrated and eluted with 10 mM Tris–HCl pH 8.0, 150 mM NaCl and 12.5 mM EDTA [25][26]. Only fractions showing the highest fluorescence yield were used within 5-7 days.

#### 2.6.2. Leakage measurements

SUVs were added into the cuvette with proteins at 0.3 mg/mL, and homogenized by rapid pipetting (about 5 sec per sample). Then, time-dependent fluorescence intensity was followed for 10 min with excitation and emission fixed at 250 and 544 nm respectively. Leakage efficiency was calculated as a percentageof the decrease in fluorescence intensity (F) respect to the initial value (Fo). The decrease of fluorescence induced by the addition of1% SDS (Fo-Ft) was taken as a 100% Leakage [25][26]. Results are shown as triplicates of 3 different experiments.

#### 2.6.3. Clearance measurement

To obtain MLV of DMPC, lipidsfrom a stock solution in chloroform were driedunder N_2_ flux as explained above, and resuspended in Tris buffer pH 7.4to a final DMPC concentration of 5 mg/mL. The tube was vortexed at room temperature for 5 min, heating at 37°C in 30-sec cycles. DMPC vesicles were added to the 0.05 mg/mL proteins samples until final molar ratio 145:1 DMPC:protein. Samples were gently mixed (for 5 sec) and clearance was determined by monitoring Optical Density (DO) at 325 nm and 24°C in a Beckman Coulter DTX 880 Microplate Reader (Beckman, CA). Curves were adjusted by fitting to a single exponential decay *y* = *y*_0_ + *ae*^−*bx*^.

### 2.7. Other analytical methods

The spatial arrangement of K107del was modeled using the SWISS-MODEL server (https://swissmodel.expasy.org, Swiss Institute of Bioinformatics Biozentrum, University of Basel Klingelbergstrasse, Basel, Switzerland) using the alignment model [27] with the apoA-I structural template obtained by Melchior et all [28]. Protein structure figures and homology models were generated using UCSF Chimera, developed by the UCSF Resource for Biocomputing [29]. For most of the experiments, protein content was quantified by the Bradford technique [30]. For the statistical analysis, datasets were analyzed in GraphPad Prism 8.0 software using unpaired parametric test. Only results with a confidence level of p <0.05 were considered. Unless otherwise stated, the results either of biophysical or biological assays were reproduced in three independent experiments and are indicated as means of triplicates ± standard deviation. Statistically significant differences between experimental conditions were evaluated by ANOVA or the Student’s test.

## 3. Results

### 3.1. Proteins Purification

ApoA-I variants (either Wt or K107del) were isolated and purified from bacterial strains in high yield and purity (Supplementary Figure 1). Previously we have shown that Wt purified under this protocol behaves almost indistinguishable from plasma apoA-I [20].

### 3.2. Leakage

In order to characterize the influence of the deletion of the positive Lys residue in position 107 on protein function, we first compared the interaction of the variant with lipid bilayers, monitoring protein-induced leakage of SUVs. Energy transfer from DPA to Tb is highly efficient in the complex confined to the vesicles internal aqueous space. If apoA-I-induced leakage occurs, excess of EDTA present in the medium replaces DPA in the complex and fluorescence decreases. As previously observed, membrane disruption mediated by Wt was smooth and slow [26] (Figure 1A and B).

**Figure 1.**
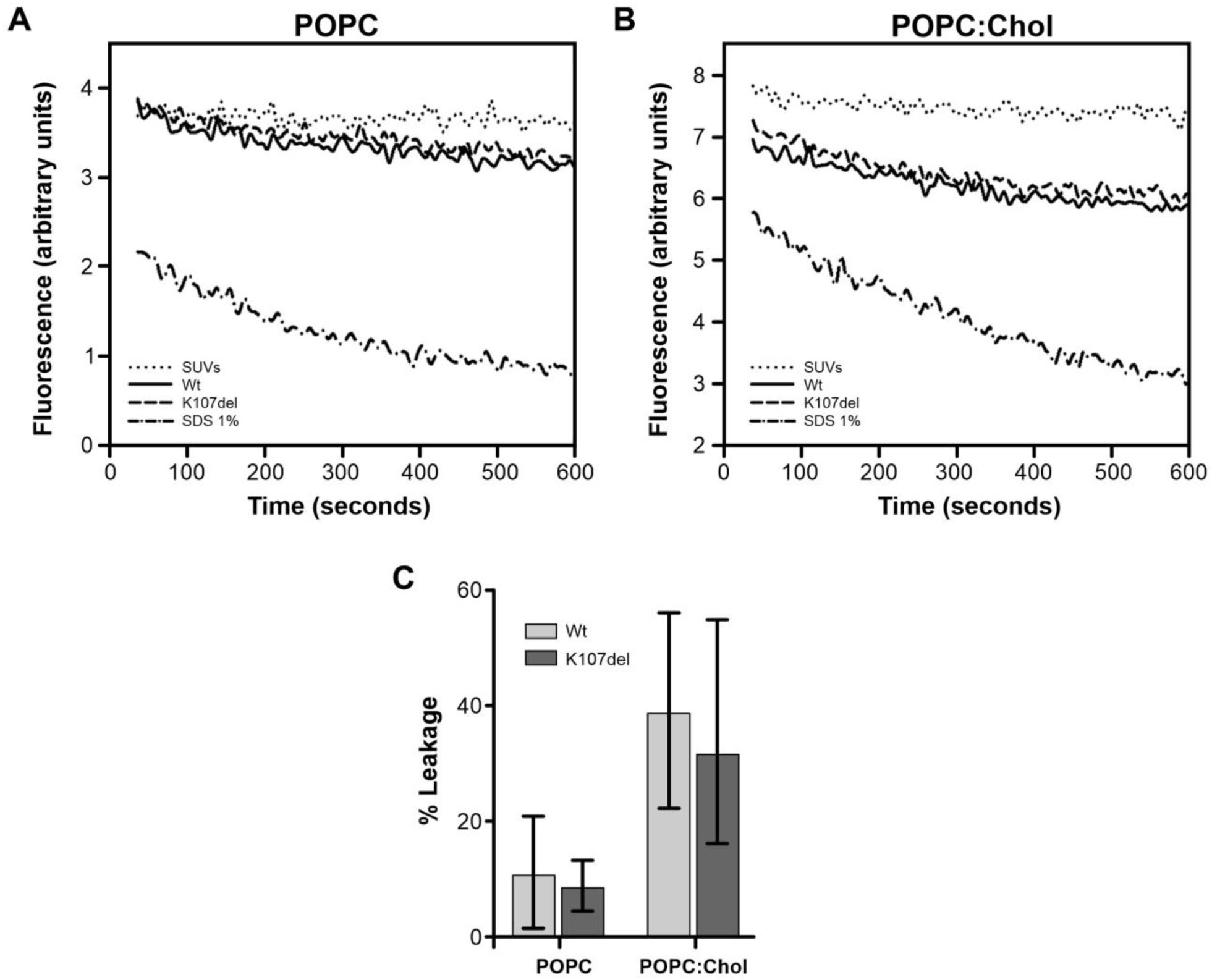
Leakage induced by Wt and K107del of Tb/DPA-loaded lipid vesicles. **A)** and **B)**, time dependence of the fluorescence intensity change obtained by adding 0.3 mg/mL of Wt or K107del to **A)** POPC or **B)** POPC:Chol SUVs. Fluorescence was registered with excitation set at 250 and emission at 544 nm. **C)** Percent of leakage was calculated to the final point (last 60 sec) after 600 sec interaction. One hundred % was taken as the decrease of fluorescence after addition of 1% SDS (dashed-point line in Figure A and B). Bars represent media ± standard deviation of triplicates of three independent measurements. No differences were found between all conditions by evaluation by t-test (p ≤0.05).

In order to compare both variants, the extension of the leakage induced by freshly folded Wt and K107del was evaluated after 10-min interaction. As Figure 1C shows, no differences were detected among both variants in their capacity to induce membrane permeation. Although a higher tendency to leakage was observed from vesicles containing Chol, the effect was neither significant nor different among variants, even at the different protein ratios tested (not shown).

### 3.3. Protein chemical treatments. Oxidation and crosslinking

#### 3.3.1. Protein structure

BS^3^ crosslinker has been widely used for the analysis of apoA-I structure [22]. Its short spacer arm (∼11 Å) makes it suitable to covalently fix primary amine groups within proteins. In addition to the N terminal amine group, apoA-I contains a high number of Lys which may be exposed to interact with the reagent in an aqueous solvent. To compare the relative efficiency of K107del to crosslink, we first diluted proteins at 0.6 mg/mL. Under those conditions, and in agreement with previous reports, Wt showed some degree of dimers covalently fixed (lane 4 in Figure 2A). This effect is less evident for the deletion mutant (lane 5). Following, we set to determine the structural effect that may occur if crosslinking is restricted to intra-chain bonds. In this trend, proteins were set at 0.05 mg/mL, as classic studies have shown mainly monomer conformation present under that concentration [22][31], and following crosslinking protein’s mobility was compared by gradient gel electrophoresis under native conditions (PAGGE). As Figure 2B shows, only monomeric species are attained under these conditions. However, the efficiency of crosslinking is evident as variants migrate slightly but significantly faster than untreated proteins, probably due to a more compact conformation attained as proteins domains are restricted by the agent.

**Figure 2.**
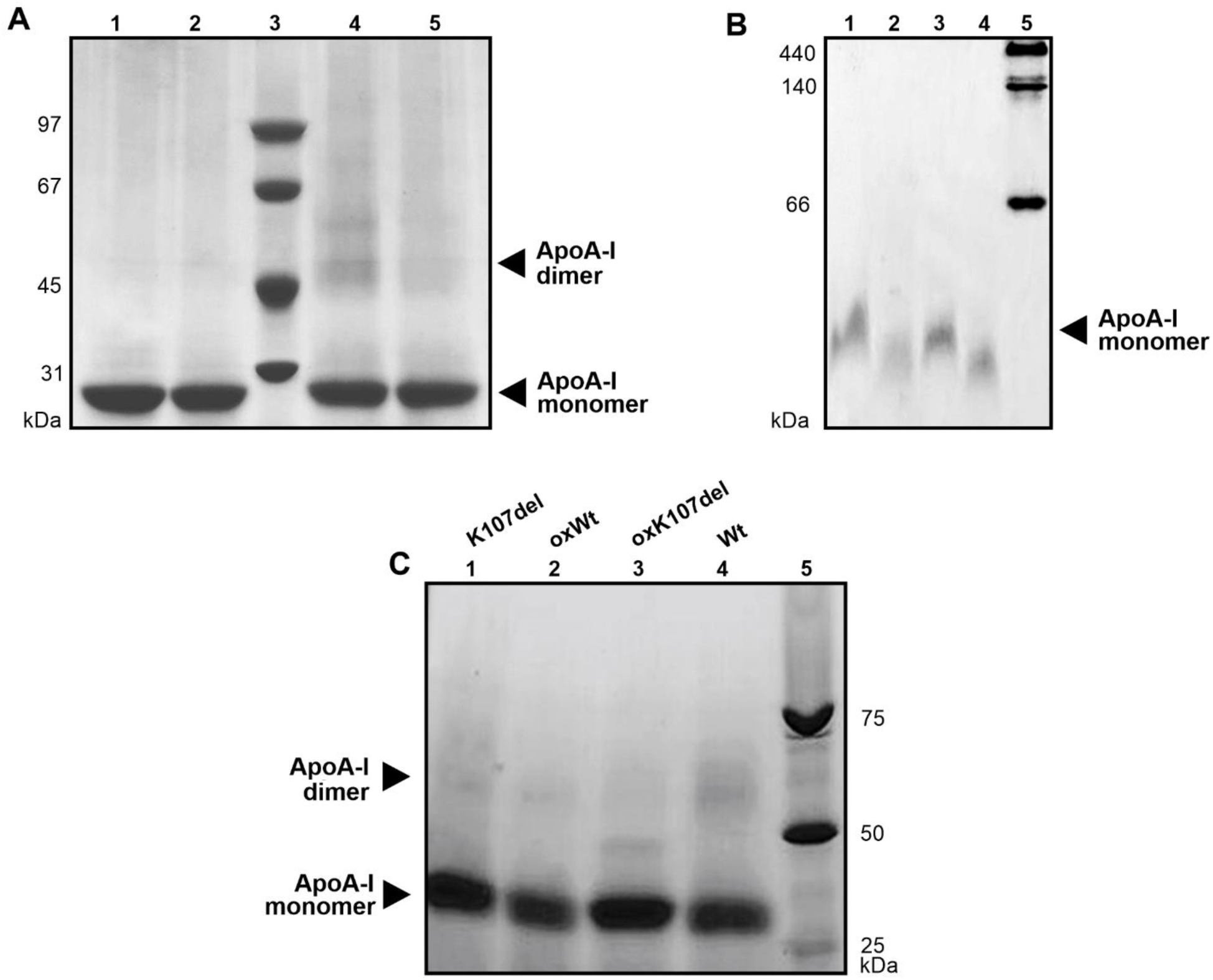
Characterization of protein cross linking by polyacrylamide gel electrophoresis. Gels were run as described and stained with Coomasie Blue. **A)** SDS PAGE at 16% of native or crosslinked proteins at 0.6 mg/mL stained with Coomasie Blue. Wt (lane 1) and K107del (lane 2) before treatment with BS^3^; lanes 4 and 5 Wt and K107del respectively after crosslinking. Lane 3 shows SDS Low molecular weight marker. Dark arrow indicates the Mw expected for the apoA-I dimer; **B)** Native PAGGE (15-20%) of proteins crosslinked at 0.05 mg/mL. Lanes 1 and 2, Wt before or after crosslinking respectively. Lanes 3 and 4, K107del before or after crosslinking respectively. Lane 5 shows commercial high molecular weight standard; **C)** SDS-PAGGE (4-16 %) to compare the effect of oxidation on the crosslinking efficiency. Lane 1 and 4 K107del and Wt treated with BS^3^ respectively. Lanes 2 and 3 correspond to oxidized Wt and K107del treated with BS^3^. Lane 5: Mw standard marker.

In order to compare whether oxidized proteins should keep the ability to self-associate, variants (either non-oxidized of after incubation with H_2_0_2_) were incubated at 0.6 mg/mL for 24 h at pH 6.0; then taken pH to 7.9, and further treated with BS^3^ as previously described (section 2.3). As it may be observed in Figure 2C, the band corresponding to dimer is vanished for the oxidized species, indicating a loss in the oligomerization capability.

The relative exposition of the Trp residues to the solvent may be used as a conformational control of protein arrangement, as it is well known that the high flexibility of apoA-I may be censored from the shift in the intrinsic fluorescence. By fixing excitation wavelength at 295 nm fluorescence is dominated in average by four Trp residues (in positions 8, 50, 72 and 108 in the native sequence) [32][33]. In agreement with previous reported results [32][34] the wavelength of maximum fluorescence of Trp (WMF) in the freshly folded K107del variant showed a small but significant 2 nm-shift to the red respect to Wt. BS^3^-crosslinking did not introduce a significant structural modification as compared with uncrosslinked neither for Wt or K107del (Figure 3A and B respectively).

**Figure 3.**
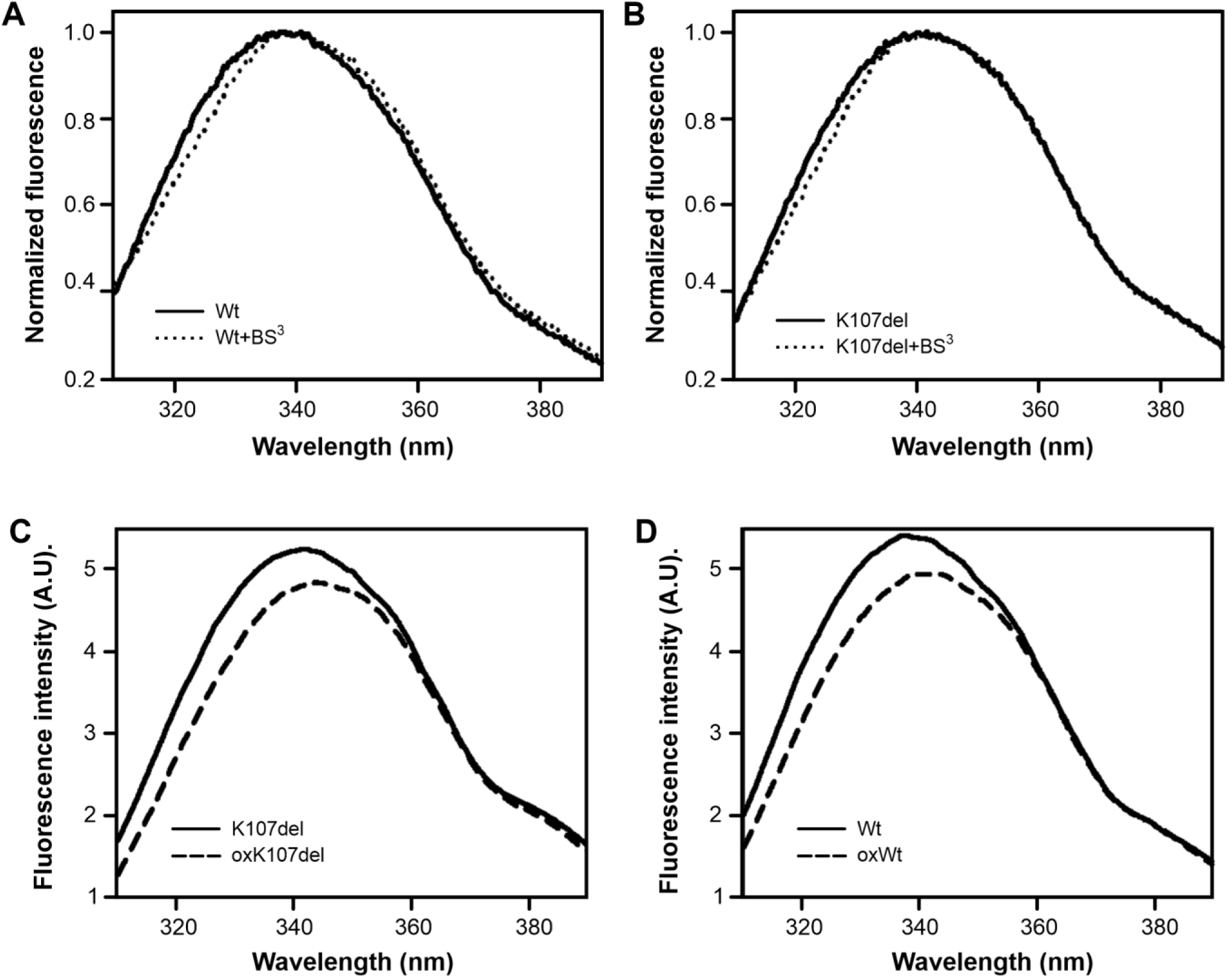
Intrinsic fluorescence spectra of posttranslational chemical-modification of apoA-I variants. Fluorescence was measured in an SLM 4800 spectrofluorometer fixing excitation wavelength at 295 nm and monitoring emission from 310 to 400 nm. **A)** Wt and **B)** K107del at 0.05 mg/mL in PBS pH 7.9 buffer either before (continuous lines) or after crosslinking (pointed lines). Spectra were normalized to the highest intensity to better compare the shift in the fluorescence emission. **C)** Wt and **D)** K107del intrinsic fluorescence registered before (continuous lines) or after oxidative treatment (dashed lines)

Following we analyzed the effect of oxidation on protein structure. We have previously described that treatment with controlled H_2_O_2_ resulted in the preservation of mainly the protein molecular weight integrity but, as expected, it induced a mild tendency to oligomerize [35][19]. To further test whether aromatic residues could be affected during the H_2_O_2_ oxidation, we checked intrinsic fluorescence of the oxidized variants. About 4 nm-shift to the red of the WMF and a small decrease in intensity indicated that the aromatic residues are relatively more exposed in both variants but significantly preserved, as extensive oxidation should result in a significant decrease in the quantum yield (Figure 3 C and D).

To better confirm the efficiency of the oxidation, we ran Mass Spec as compared with the fresh variants. As it was previously reported, this treatment should oxidize the three Met present in the Wt (86, 112 and 148 residues) which might result in a gain in 48 Da. The Mw obtained for oxWt (Figure 4A, 27,857 Da) or oxK107del (Figure 4B, 27,727 Da) were each 71 Da-higher than the values obtained for the native variants [19], which supported a complete oxidation of Met residues. The excess respect to the expected value (23 Da) should be explained by ionic adducts from sodium (already described for apoA-I [36]). In addition of a main peak corresponding to the monomer, a small peak of 56 kDa may indicate small amount of Wt dimer present in the sample (inset Figure 4A).

**Figure 4.**
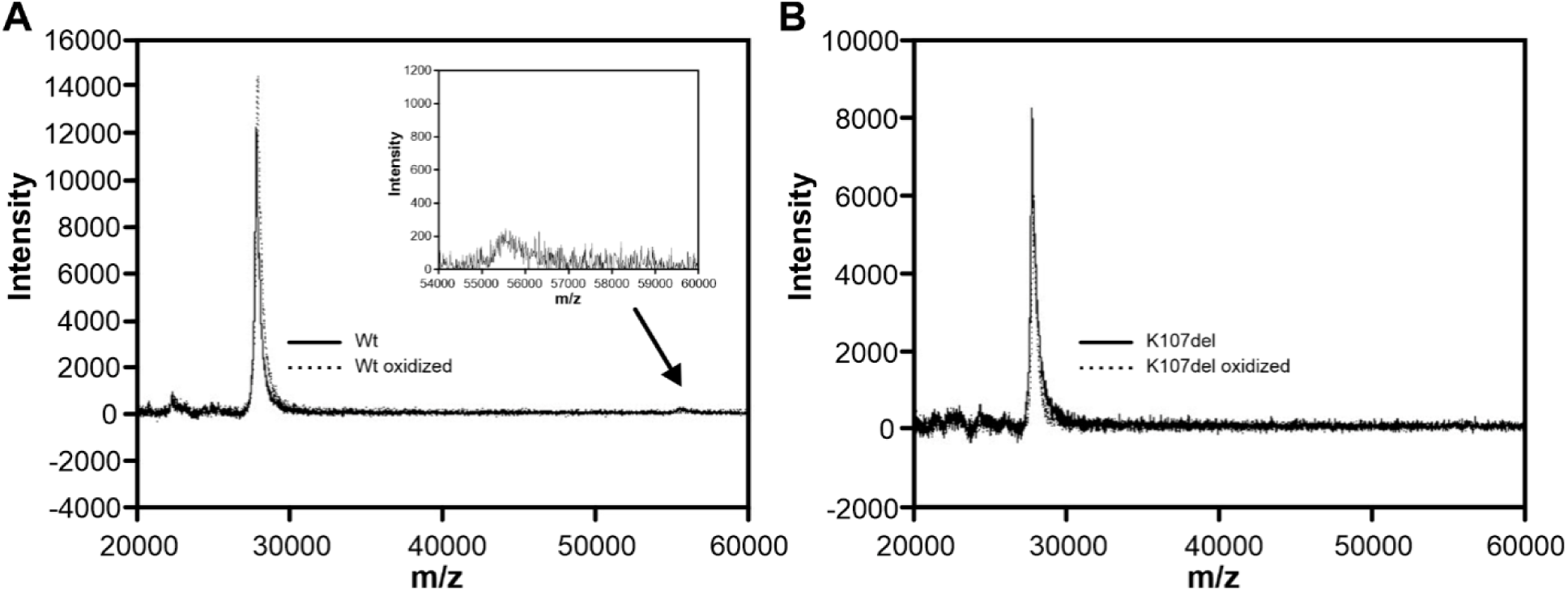
Effect of oxidation on proteins molecular weight. MS spectra of **A)** Wt and oxWt and **B)** K107del and oxK107del. Inset in Figure A) amplification of the peak shown by the black arrow, indicative of dimeric apoA-I

#### 3.3.2. Effect of chemical modification on protein function

With the aim to answer whether chemical modifications may affect proteins function, we performed a well-established test, as it is the ability of apoA-I to solubilize lipids from vesicles at their transition temperature [33]. Figure 5A shows that, in agreement with the leakage experiments, the deletion of Lys107did not modify its efficiency to solubilize lipids. Chemical modification of proteins by oxidation reflects a similar result showing no significant differences to solubilize lipids between the oxWt and oxK107del. However oxidized proteins presented an increased capacity to clarify MLV liposomes respect to the native protein conformations.Evaluation of the structural restriction induced by intramolecular crosslinking resulted in drastic impairment of the variants’ ability to induce clearance, being more affected Wt than K107del. Significant differences in the clearance at the final point are compared in Figure 5B.

**Figure 5:**
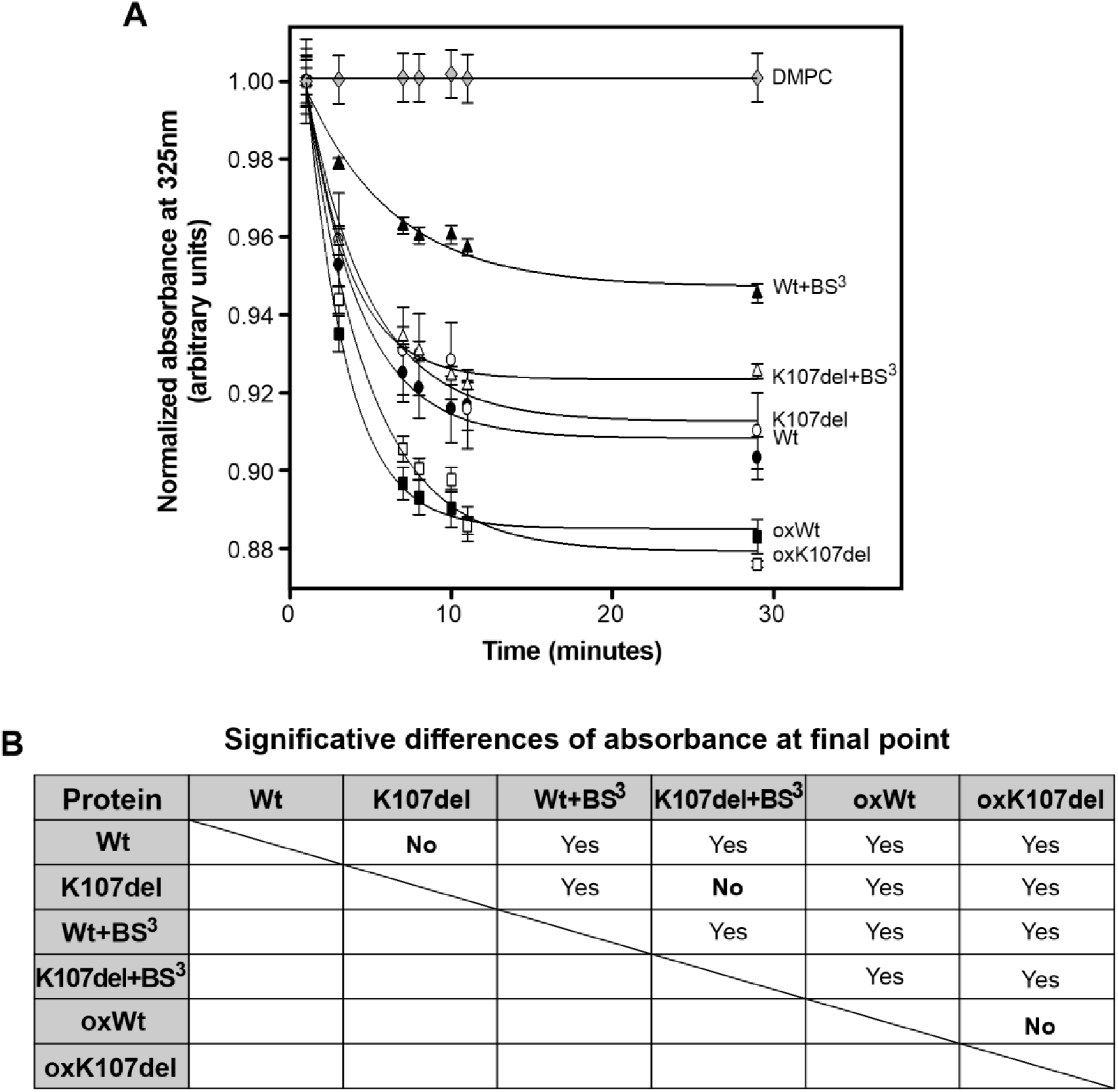
Time-course of the interaction of native, crosslinked or oxidized variants with DMPC liposomes. **A)** Multilamellar DMPC vesicles were added to apoA-I variants at 0.05 mg/mL, to a final molar ratio of 145:1 phospholipid to protein at 24 °C. Lipid solubilization efficiency was measured by following Absorbance at 325 nm. Continuous lines correspond to the fitting of the curves to single exponential decay. **B)** Difference among the sample’s absorbance at the last incubation time (30 min) was estimated by the t-test (p≤ 0.05)

Finally, and in order to better estimate a possible explanation for the behavior of K107del respect to Wt, we built a model of the variant using as a template the recent consensus structure of the monomeric human apoA-I proposed from combined biophysical evidences [28]. As Figure 6 indicates, our modeling predicts a distortion of helix 4 due to the deletion, which may weaken inter chain salt bridge interactions between neighbor Glutamate and Lysine residues. The estimation of the solvent accessible surface area (SASA) was performed by the PyMOL Molecular Graphics System, Version 2.0 (Schrodinger, LLC), indicating a relative increase of the area exposed for K107del (15,936 Å^2^) with respect to the Wt (15,651 Å^2^).

**Figure 6:**
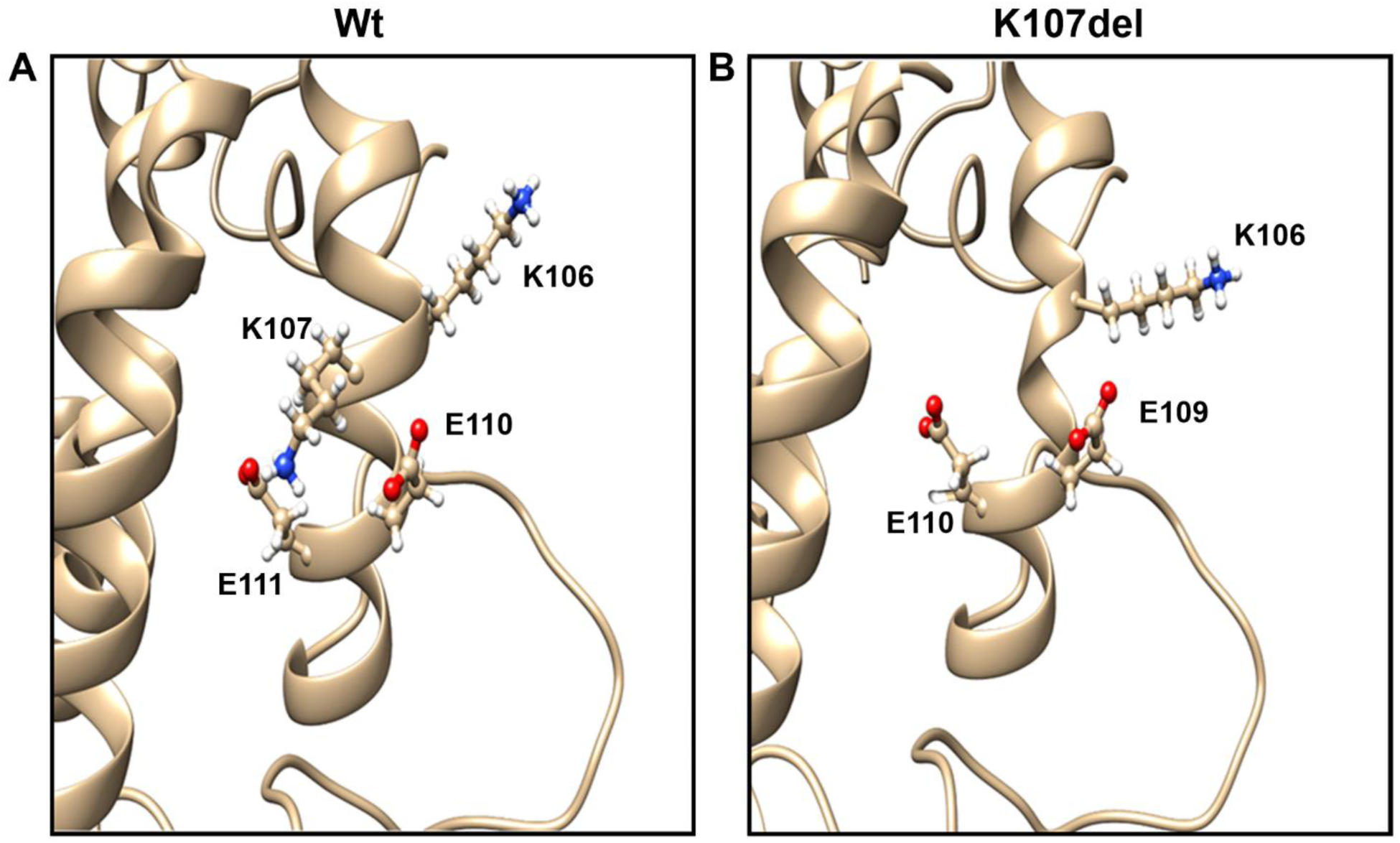
Model of the apoA-I and K107del arrangement. Crystallographic structure of **A)** apoA-I Wt or**B)** K107del modeled using the SWISS-MODEL server (https://swissmodel.expasy.org), with the apoA-I structural template obtained by Melchior et all [28]. Protein main sequence is shown in ribbon diagram. Both images show the amino acids lysine and glutamate in the form of balls and sticks.

## 4. Discussion and conclusions

In the present study, we set out to characterize local conditions that may shift apoA-I structure from the native folding, inducing its aggregation and dysfunction in the atherosclerotic plaques. From the analysis of this chronic pro inflammatory scenario, common factors are to be considered: 1) chemical modifications due to ROS freed by activated macrophages, among those oxidation and crosslinking [35]; 2) intrinsic modifications caused by hereditary mutations in the apoA-I sequence which should turn it prone to misfold or misfunction. In this trend the deletion mutant K107del was identified as a good target, as it was associated with severe atherosclerosis in patients [37][38]. Studies done either in *in vitro* or *in vivo* models have not completely determinedneither the reasons for the failure of apoA-I to fulfill the atheroprotective role nor the increment in the severity of the lesions associated with the deletion variant.

Although lower-than-normal levels of HDL were usually attributed to the phenotype of patients carrying apoA-I K107del [37][38][39] small clinical and biophysical differences respect to the native protein were reported, and sometimes with discrepancies among the assays. While Rall et al found reduced efficiency to activate LCAT of isolated K107del *in vitro* [40], this function was shown similar to Wt in other studies [41][42]. In a similar trend, lipid binding affinity was reported either decreased [41], or similar to Wt [43]. Cholesterol efflux from cultured cells was not significantly altered by the deletion [44][45]. In other patients this mutant was associated withhypertriglyceridemia [39], and *in vitro* studies supported an increased binding to triglyceride-rich particles [43]. These findings suggest that, in addition to the lower efficiency that the mutation may induce in the normal apoA-I functions, the effect does not seem to be drastic. Instead, the Wt protein that is also present in heterozygote subjects, with a dissimilar prevalence of the mutated over the normal allele in all affected individuals may counterbalance minor shifts, which could explain -at least in part-the variety in the observationson patients.

Here we first tested the hypothesis that the deletion of the positive residue in position 107 induces a structural shift affecting proteins folding and function. In concordance with previous reports [32][43] our data support a more flexible structure induced by the deletion. The lower efficiency of inter-chain crosslinkingat concentrations greater than 0.1 mg/mL (Figure 2A), may indicate a tridimensional protein folding that is more relaxed than the Wt. In agreement with this, a small amount of Wt dimer could be detected in the Mass Spec analysis (inset in Figure 4A) which is absent for the deletion mutant.The small red shift in the intrinsic Trp fluorescence of K107del shown here, and the increased binding to ANS and Bis-ANS probes previously reported [19], also support a more flexible structure of the monomeric conformation.

Based on the class A model of alpha helices, Lys 107 should occur in a segment of the repeating 22-residue amphiphilic helices [46]. The traditional helix wheel simulations predict that the deletion of this residue induces a ∼90° shift of helix axe, disrupting the nature and orientation of the hydrophobic face [40]. More recently, analysis based on the reported crystallography structure of the C-terminal truncated apoA-I [47], suggested that this mutation induces a weakening of the salt bridge networks [43]. To better strength this observation, we modeled and compared Wt and K107del by the SWISS MODEL predictor. As Figure 6 suggests, a less compact domain might be generated due to the absence of that residue. Whether one or all the models predict the effect of the mutation is not known and must be further studied.

In spite of the observed structural shift, neither lipid solubilization nor leakage of vesicles is significantly affected by the mutation, which indicates that some other features may contribute to explain the strong clinical manifestations detected in patients.The finding of dysfunctional HDL and apoA-I in the atherosclerosis plaque suggests that the protein may be a *victim* of the pro-inflammatory scenario. As we show here, lipid function under intramolecular crosslinking is impaired being Wt more restrained from lipid solubilization than K107del (Figure 5). In concordance with previous reports [32], the higher flexibility of K107del lipid-free protein may allow its reorganization as recruiting lipids even under some structural restraints following crosslinking. It remains to be determined if this effect on K107del is caused by crosslinking of non-crucial residues involved in interaction with lipids domains or due to major flexibility of mutant allows to uncrosslink flexible-key domains (helices) to form the belt in discoidal HDL. In order to confirm this, specific experiments remain to be done reviewing the role of the amino group in position 107.

In addition to croslinking, other critical post translational modification is worth to be considered such as oxidation.In this trend we first set to analyze the probable loss in the efficiency of lipid solubilization. Instead, an increase in lipid binding affinity is observed as Wt (and similarly is observed for K107del) is oxidized under our experimental conditions, which is in agreement with Panzenböck observations on Wt with Met86 and Met112 oxidized [48]. The reasons of this behavior is not known but, as they suggest, it may be hypothesized that the introduction of a negative charge (sulfoxide groups) in the place of non polar Met at the boundary between a polar and non polar faces may alter the lipid binding affinity [48]. In this trend, we have recently reported a similar effect behavior as the non polar Leu was substituted by a polar Arg [49].

In addition to a loss of function, the effect of the environment on the tendency of the proteins to aggregate is worth to be analyzed. The participation of apoA-I in age-related atherosclerosis in association with amyloidosis was previously raised [15][16][21]. Aortic intimaamyloid deposits are often associated with atherosclerosis and apoA-I is usually described as diffuse patches within these lesions [15]. The presence of K107del inducing amyloidosis associated with severe atherosclerosis may indicate that the mutation has a strong propensity to participate in the crosstalk among these chronic pathologies. As oxidation is other event proposed to be prevalent in inflammatory scenarios we have considered this possibility as a chemical modification. A loss in the capacity to acquire a functional dimer conformation (as shown in Figure 2C) or a higher tendency to aggregate of the oxidized variant may induce not only the impairment in protein function but in addition to yield a citotoxic amyloid prone species that could worsen the pro-inflammatory scenario. In this regard, we have recently shown that oxK107del is more sensitive than oxWt to yield amyloid fibrils [19]. In addition to K107del, another mutant L178P was described associated with severe atherosclerosis [34]. Interestingly this variant was also reported within a laryngeal presentation of systemic apolipoprotein A-I-derived amyloidosis, which might strength the close association between both chronic pathologies [44]. The correlation of L178P with intimal amyloidosis was not reported, but it is not clear whether it was analyzed as it might not be discarded.

As a conclusion, we suggest that it is required drastic protein processing within the microenvironment, as oxidation and crosslinkg, probably more than one synergizing, to induce a collapse of protein structure which is associate to both, protein loss-of-function (as lipid-binding), and aggregation as amyloid complexes.

## Acknowledgements

This work was supported by the Consejo Nacional de Investigaciones Científicas y Técnicas (CONICET, PUE 22920160100002 to HG); Agencia Nacional de Promoción Científica y Tecnológica (ANPCyT, PICT-2016-0849 to MAT and PICT-2016-0915 to HG); Universidad Nacional de La Plata (UNLP) (M187 to MAT).

## Author contribution statement

RAG and IDL performed the experiments, did data analysis and designed the research; HAG did data analysis and searched for funding, MCG did data analysis and discussed results, NAR and MAT designed the research, wrote the manuscript and searched for funding.

