## Supplementary Figure 1 for "Understanding the role of apolipoproteinA-I in atherosclerosis. Post-translational modifications synergize dysfunction?"

### Supplemental Figure 1

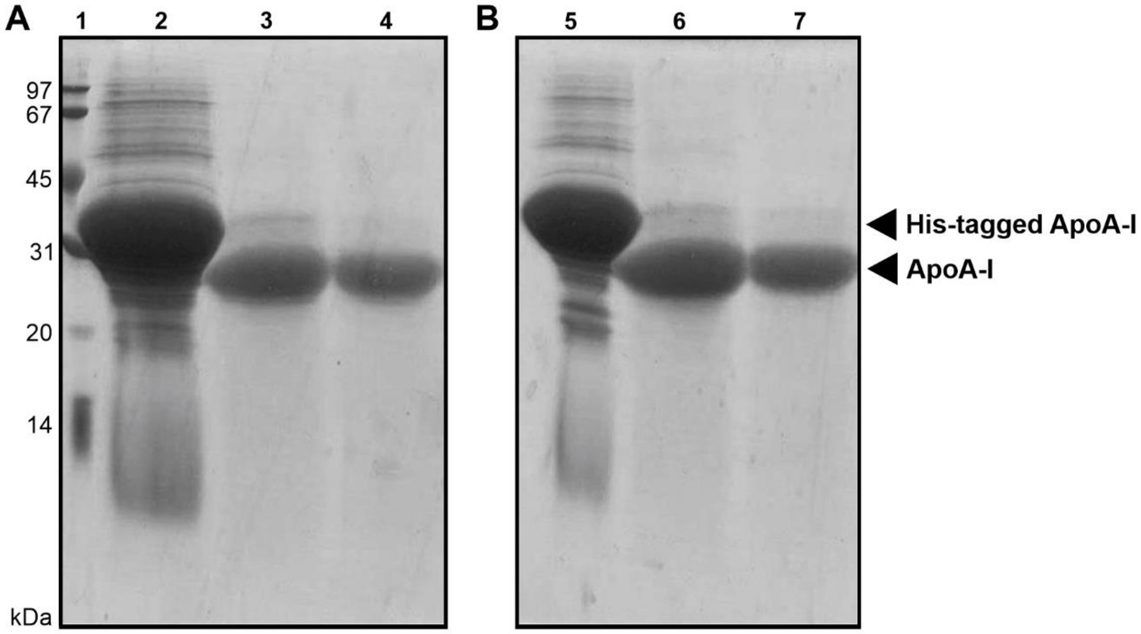


**Figure 1 Characterization of protein purity.** K107del (A) and Wt (B) were purified by IMAC Sepharose affinity columns and loaded onto 16 % SDS-PAGE stained with Coomasie Blue. Lane 1: commercial low molecular weight standard LMW standard (GE Healthcare). Lanes 2 and 5: His-tagged protein. Lanes 3 and 6: final pure fractions at 0.9 mg/mL. Lanes 4 and 7: final pure fractions at 0.5 mg/mL.
